# A Multi-Species Functional Embedding Integrating Sequence and Network Structure

**DOI:** 10.1101/229211

**Authors:** Jason Fan, Anthony Cannistra, Inbar Fried, Tim Lim, Thomas Schaffner, Mark Crovella, Benjamin Hescott, Mark D.M. Leiserson

## Abstract

A key challenge to transferring knowledge between species is that different species have fundamentally different genetic architectures. Initial computational approaches to transfer knowledge across species have relied on measures of heredity such as genetic homology, but these approaches suffer from limitations. First, only a small subset of genes have homologs, limiting the amount of knowledge that can be transferred, and second, genes change or repurpose functions, complicating the transfer of knowledge. Many approaches address this problem by expanding the notion of homology by leveraging high-throughput genomic and proteomic measurements, such as through network alignment.

In this work, we take a new approach to transferring knowledge across species by expanding the notion of homology through explicit measures of functional similarity between proteins in different species. Specifically, our kernel-based method, Handl *(Homology Assessment across Networks using Diffusion and Landmarks)*, integrates sequence and network structure to create a *functional embedding* in which proteins from different species are embedded in the same vector space. We show that inner products in this space capture functional similarity across species, and the vectors themselves are useful for a variety of cross species tasks. We perform the first whole-genome method for predicting phenologs, generating many that were previously identified, but also predicting new phenologs supported from the biological literature. We also demonstrate the Handl-embedding captures pairwise gene function, in that gene pairs with synthetic lethal interactions are co-located in Handl-space both within and across species. Software for the Handl algorithm is available at http://github.com/lrgr/HANDL.

## 1 Introduction

A primary challenge of research with model organisms is to transfer knowledge of genetics – i.e. a mapping of genotype to phenotype – between model organisms and humans. The main promise of researching model organisms stems from researchers’ ability to measure the organisms in ways that are infeasible in humans. To realize the promise of this research, it is crucial to transfer knowledge between species – ideally, in two directions. First, discoveries in model organisms can be transferred to improve knowledge of human genetics (e.g. via homology^1^). Second, knowledge of human genetics can be transferred to design better experiments in model organisms (e.g. for disease models).

More specifically, cross-species knowledge transfer can enable a wide variety of applications. First and foremost has been the large-scale annotation of protein function by transferring function annotations (e.g. from the Gene Ontology [4]). Addressing this problem remains valuable, even in the era of high-throughput genomics, as fewer than 1% of protein sequences in Uniprot have experimentally-derived functional annotations [63]. A second but less well-explored application is in predicting human disease models through ‘orthologous phenotypes’ or *phenologs* [38]. McGary, et al. [38] reasoned that while conserved genes may retain their *molecular* functions across species, conserved molecular function may manifest as different “species-level” phenotypes. As such, they introduced a statistical test to identify such phenologs. Another application of cross-species knowledge transfer is for *pairwise* gene function (*genetic interactions*). Knowledge of synthetic lethal genetic interactions is crucial for the study of functional genomics and disease [45,46]. Since measuring synthetic lethal interactions in humans is currently infeasible, computationally transferring knowledge of these interactions from model organisms (such as yeast or mouse) to humans (and human cancers) has become a focus of recent research.

Nonetheless, cross-species knowledge transfer remain quite challenging because many model organisms diverged from humans millions of years ago and have fundamentally different genetic architectures. Initial computational approaches to transfer knowledge across species relied on measures of genetic heredity such as *homology*. Using genetic heredity to characterize genes in different species is foundational to genetics and comparative genomics as there is widespread and long-standing evidence that conserved genes tend to share the same function [32]. Consequently, methods for inferring homology are crucial for transferring knowledge prompting researchers to develop sophisticated algorithms to infer homology from DNA or protein sequences (e.g. [47]), and protein-protein interaction (PPI) networks.

The most common method to infer homology is the Basic Local Alignment Search Tool (BLAST) which computes a sequence similarity score between pairs of proteins (or genes) in different species, and reports the statistically significant pairs as homologs [3,57]. A second class of methods expands beyond sequence by using protein structure to infer remote homology with the goal of classification and function prediction [23].

Despite their widespread use, current approaches that infer homology from sequence data face several challenges that limit the amount of knowledge that can be transferred. Structure-based methods are computationally expensive and do not currently scale to entire genomes. Moreover, in many cases, only a relatively small subset of genes between species have sequence homologs. Further, as species diverge, protein functions change and are re-purposed (e.g. [21]), and genetic interactions are often rewired [56,62]. Thus, more recent approaches aim to expand the notion of homology to capture both convergent (analogous) and divergent (homologous) evolutionary mechanisms.

Recent methods expand beyond sequence homology by using widely-available high-throughput proteomic datasets. Many of these methods use genome-scale protein-protein interaction (PPI) networks, as genes with similar functions tend to have similar topology (e.g. as measured by network propagation [14]). The comparison of PPI networks across species is well studied and commonly referred to as the network alignment problem [36,50,53,30,39,37,1,65,28,59,41,25]. The goal of network alignment is to establish a mapping between nodes in different networks. The dominant paradigm for this work is based on matching nodes; in some cases, this is done using cross-species node similarity scores in an intermediate step (e.g., [58,36,39,37,44,24]). Working in another direction, Jacunski, et al. [26] introduced the notion of *connectivity homology* to relate genes in different species by their position in corresponding PPI networks. While not the focus of their paper, they show that pairs of genes with the same or similar functions have lower connectivity homology dissimilarity scores. More recently, Khurana, et al. [29] developed an embedding for proteins in multiple species for an application concerning neurodegenerative diseases.

### 1.1 Contributions

In this work, we go beyond matching-oriented network alignment and generalize the notion of similarity scores by creating a biologically meaningful vector space into which we embed the nodes of multiple networks. Both the similarity scores derived from this embedding as well as the node vectors themselves are useful for a variety of tasks.

Our method makes only two assumptions. First, it assumes that network homology can be captured using a similarity function that is a *kernel*, which encompasses a broad class of useful metrics. Second, it assumes that sequence homology is known for some subset of proteins across the different species. We illustrate the method in this paper using pairs of species, but extension to simultaneous treatment of multiple species is straightforward.

We note that network alignment methods do in some cases compute similarity scores internally. However, such methods construct these scores only for the purpose of node matching, limiting their applicability beyond alignment and protein function annotation. It is not obvious how to extend these methods for problems that go beyond node matching – such as transferring knowledge of *pairwise* gene function (e.g. synthetic lethality). More recently, Jacunski, et al. [26] introduced a general purpose cross-species node vector representation, but their work is focused on training classifiers using the vectors as features. Although they show good performance on these prediction tasks, their representations are derived from summary statistics, and we show the distances in their vector space are not strongly correlated with functional similarity.

In this paper, we use a *diffusion* kernel to capture functional similarity and call the resulting method Handl *(Homology Assessment across Networks using Diffusion and Landmarks)*. Diffusion kernels are a natural tool for capturing aspects of local and global network structure that correlate with functional similarity of nodes. Our method may be used with other kernels to relate proteins for other biological applications, or to assess other network properties.

We demonstrate that Handl homology scores capture functional similarity between proteins in different species, benchmark Handl against two standard network alignment algorithms and connectivity homology, and find that Handl performs as well as or better than these other methods, depending on the evaluation metric. We show that comparisons in the Handl vector space can replace standard sequence homology for two important applications:

1. **Phenologs.** We show the first whole-genome method for predicting phenologs. The method considers sets of phenotype-associated genes from two species and evaluates their orthology using the statistical test from [38]. In contrast to [38], we do not only consider known homologs in the statistical test; instead we consider *any* protein pairs that Handl predicts are functionally close across both genomes. A subset of our predicted phenologs match those identified in [38]; additionally, we also predict many new phenologs and support these new predictions with biological literature.
2. **Cross-species synthetic lethality.** We demonstrate that Handl-embeddings for *pairs* of genes in different species capture pairwise gene function in *S. cerevisiae* (*S.c.*) and *S. pombe* (*S.p.*). We show that gene pairs with synthetic lethal (SL) interactions from both species are co-located in Handl-space. Using Handl-embeddings as features, we find that classifiers can learn to identify SLs in multiple species *simultaneously*, achieving an area under the receiver-operator characteristic curve (AUROC) of at least 0.88 on held-out data in both *S.c.* and *S.p.* The classifiers using Handl-embeddings as features outperform SINaTRA on 7 of 12 evaluation metrics on the datasets.

Together, these tasks encompass transferring knowledge both between model organisms, and between model organisms and humans. We conclude with a proof-of-concept experiment using Handl to predict protein function in humans, simultaneously leveraging networks from humans and multiple model organisms.

## 2 Results

### 2.1 Handl algorithm

The central contribution we make in this paper is to introduce a *network-based metric of functional similarity between proteins of different species.* In this section we introduce and motivate our metric.

Handl, as shown in Figure 1, leverages properties of kernel functions as tools for measuring the similarity of nodes in a network. While the use of kernels for the study of individual networks is well known [14], it remains an open problem to construct network-based kernels that capture the similarity of nodes between *different* networks. This is the challenge that Handl addresses.

**Fig. 1:**
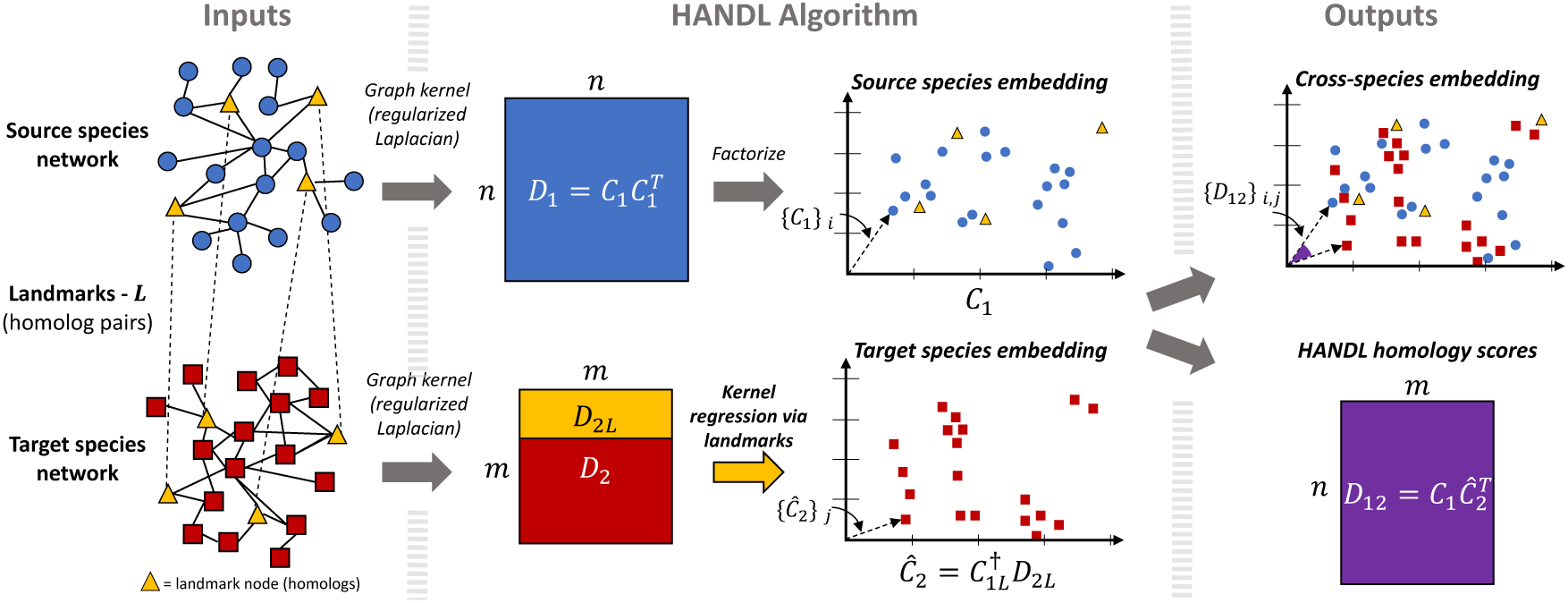
Given a source PPI network, a target PPI network, and a set of landmark (homolog) pairs across species, Handl computes diffusion kernels for each network. Then, Handl factorizes the diffusion kernel for the source species into its reproducing kernel Hilbert space (RKHS). Finally, Handl solves a linear system of the source species’ RKHS and the target species’ diffusion kernel to create a cross-species vector embedding of source and target proteins. The inner products of these embeddings correlate with functional similarities and the embeddings themselves allow for functional comparisons between proteins across the two networks.

Starting from a given kernel (node similarity function), and given the PPI networks for a source and a target species, Handl starts by performing a kernel embedding of the source species nodes (proteins). That is, for each node *v*_*i*_ in the source species, Handl computes a vector *ϕ*(*v*_*i*_) such that the vector inner product for any two nodes is equal to the kernel similarity of the nodes. The vectors *ϕ*(*v*_*i*_) can be thought of as an embedding of the source network into a geometric space.

The key step in Handl is to also embed the nodes of the target species in the *same* vector space. Handl does this through the use of landmarks – nodes that are known to be the same in both species. For PPI networks, homologous proteins play the role of landmarks. Handl then places target nodes in the vector space so as to capture their similarity to the landmark nodes.

The result is a joint embedding of both networks in the same vector space. Because both source and target nodes are placed in a way that captures similarity to the same set of landmarks, it becomes possible to score and estimate similarities between nodes in different species. We refer to these cross-species similarity scores as Handl *homology scores*, the space of the embedding as Handl*-space*, and the set of vectors for each network as Handl*-embeddings.* It is also useful to create embeddings for pairs of proteins, in which case we simply add together the embeddings of two proteins. More details and rationale are provided in Section 4.

While Handl can be used with any network kernel, in this study, we use the regularized Laplacian network kernel in order to capture functional similarity between proteins in different species. The regularized Laplacian kernel is a natural choice for this task because of its close relationship to the principle of “guilt-by-association” often used by protein function prediction methods [35], and to network diffusion methods (e.g. see [64]). For additional motivation and details of our choice of kernel, see Section 4.2.

Our results show that the Handl-embeddings encode functional relationships between proteins in different species. Leveraging Handl homology scores as well as Handl-embeddings, we successfully perform three cross-species tasks that traditionally use homologs: protein function annotation, phenolog discovery, and synthetic lethal classification.

### 2.2 Handl-embeddings of protein-protein interaction networks

For the experiments in this study, we compute the regularized Laplacian for human (*Homo sapiens*, or *H.s.*), mouse (*Mus musculus*, or *M.m.*), baker’s yeast (*S. cerevisiae*, or *S.c.*) and fission yeast (*S. pombe*, or *S.p.*) PPI networks. We then construct Handl-embeddings for human (source) - mouse (target), human-*S.c.*, mouse-*S.c.* and *S.c.*-*S.p.* We downloaded and processed PPI networks for human and mouse from the STRING database [20], PPI networks for *S.c.* and *S.p.* from BioGRID database [60], and lists of homologs from Homologene [57]. We restrict our analysis to the two-core of the largest connected component of each network. After processing, the networks range in size from 1,865 nodes (*S.p.*) to 12,872 nodes (human). For each pair of species, we used 400 randomly selected pairs of homologs to compute Handl-embeddings and Handl homology scores. See Section 4.4 for additional details on data sources and processing.

We find that Handl-embeddings for networks of various sizes can be computed in a practical amount of time. Using an Intel Xeon E6-2660 v2 processor with 20 hyper-threaded cores (40 threads) and 94GB of memory, Handl-embeddings from human to mouse, mouse to human, *S.c.* to *S.p.*, and *S.p.* to *S.c.*, can be computed in 249.1, 3.5, 4.1, and 0.25 minutes, respectively.

### 2.3 Handl homology scores capture functional similarity across species

Our results show that high Handl homology scores are strongly correlated with functional similarity between human and mouse proteins. For this section we evaluate results for pairs of proteins (*p*_*i*_, *p*_*j*_), where *p*_*i*_ and *p_j_* are from human (source) and mouse (target), respectively. We only include pairs for which neither *p*_*i*_ or *p*_*j*_ are part of a landmark pair used to compute Handl embeddings and Handl homology scores.

We use the Resnik score [54] as a quantitative measure of functional similarity. The Resnik score between two Gene Ontology (GO) [4] terms is the information content of their most informative common ancestor in the GO hierarchy; to compare two proteins we take the maximum Resnik score over all pairs of GO terms. The Resnik score has been shown to be one of the best performing metrics for capturing functional similarity within the GO hierarchy [51].

To demonstrate the relation between Handl homology score and functional similarity, we order each pair according to their Handl homology scores, and plot rankings against the Resnik score of the pair. The results (smoothed over non-overlapping windows of 100,000 observations) are shown in Figure 2a. Other cases are shown in Supplemental Information S3.1.

**Fig. 2:**
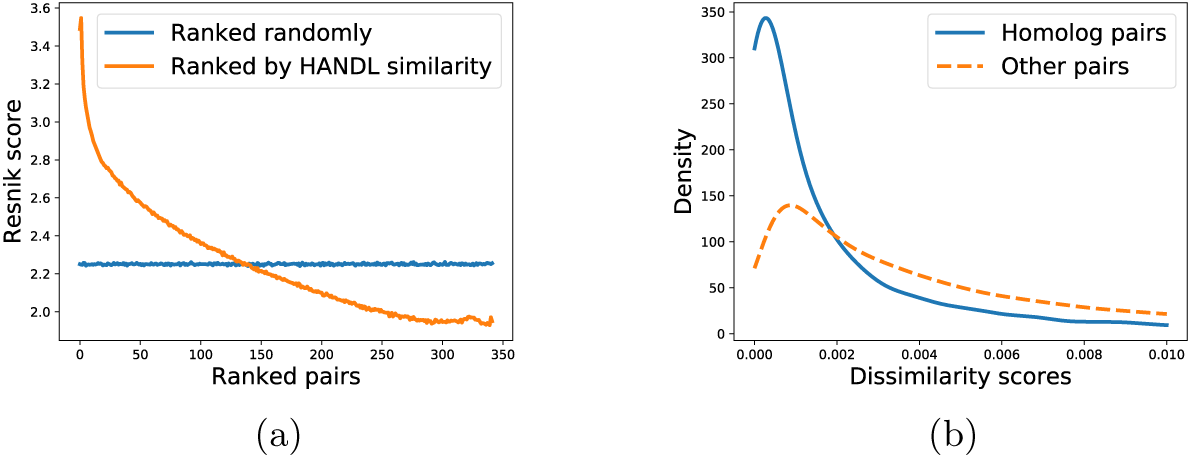
The relationship between Handl homology scores and functional similarity for the human (source) to mouse (target) embeddings. (a) Relationship between functional similarity measured by Resnik score (y-axis) and protein pairs ranked (x-axis) by Handl similarity (shown in orange) and ranked randomly (shown in blue; included as a baseline). (b) Distribution of Handl dissimilarity scores for homologous protein pairs compared to distribution for other (non-homologous) protein pairs.

The figure shows that Handl homology scores are strongly correlated with functional similarity across the entire range of scores. Furthermore, the very largest Handl homology scores are indicative of protein pairs with particularly high functional similarity.

#### Homolog pairs have distinct Handl homology scores

Next, we show that pairs that are known to be functionally related are distinguishable by their Handl homology scores. For this purpose, we separate pairs (*p*_*i*_, *p*_*j*_) where *p*_*i*_ and *p*_*j*_ are homologous proteins in different organisms from other pairs and compare their Handl homology scores.

In Figure 2b, we show the distribution of Handl dissimilarity among known homolog pairs, as compared to the distribution of scores across other pairs. In this figure, we use reciprocal scores (dissimilarities), meaning that small scores are associated with high functional similarity. Only the left side of the distributions are shown, as the distribution of all pairs extends far to the right and obscures the homolog distribution on the left. As suggested by the plots, the mean Handl homology scores for human-mouse homologs are 36% lower than the mean Handl homology scores across other protein pairs. Other cases are shown in Supplemental Information S3.1. These results provide additional evidence that Handl homology scores are correlated with functional similarity across species.

#### Handl captures shared biological information beyond node degree

Furthermore, we find that Handl captures shared biological information from network topology and not only from node degrees. We assess the statistical significance of the difference in Handl homology scores between homologous and non-homologous pairs by generating 1000 pairs of random networks in which each node is given degree very close to that in the original network, but in which edges have been randomized. Specifically, we follow the method of Newman et. al [43] to generate graphs with given degree distributions and remove self loops and parallel edges afterwards. We then compute an empirical *P*-value by counting the number of pairs of random networks for which the difference in mean Handl homology scores between homologs and non-homologs is greater than that observed in real PPI networks. Given the expense of generating many permutations of the large human network, we instead assess the two yeast (*S.c.* and *S.p.*) networks. We find that Handl-embeddings for homologous pairs have statistically significantly higher Handl homology scores (*P* = 0.002 for embedding *S.c.* to *S.p.*, and *P* = 0.005 for embedding *S.p.* to *S.c.*). Consequently, we conclude that the differences in the distributions of Handl homology scores for homologs and non-homologs is unlikely to be due to node degree alone, but instead is a result of Handl capturing more detailed network topology.

#### Comparing Handl to other methods

We compare Handl to network alignment and other algorithms in its ability predict cross-species functional similarity of proteins. In particular, we compare Handl to two standard network alignment algorithms, IsoRank and HubAlign [36,24], from which it is possible to extract cross-species protein metrics that can be interpreted as similarity scores, and thus serve as a natural comparison point for Handl homology scores. We also compare Handl to the sequence alignment algorithm BLAST [3] and SINaTRA [26].

We then define a pair of proteins (*p*_*i*_, *p*_*j*_) from two species to be *k-functionally similar* if both *p*_*i*_ and *p*_*j*_ are annotated by the same GO term and, in each species, that GO term is associated with at most *k* proteins (see Section 4.4 for details on processing of GO). This metric for functional similarity was also used (without naming it) in [26] and is similar to the Gene Ontology Consistency (GOC), or Functional Consistency (FC) metrics that are also used to evaluate network alignment algorithms [12,24].

We then rank cross-species protein pairs by similarity scores obtained from Handl and other benchmarked algorithms and compute enrichment of *k*-functional similar pairs at *k* = 100 in the top ranked sets. For Handl, we rank pairs by Handl homology score. For IsoRank and HubAlign, we rank pairs by similarity scores from the scoring matrix used to generate network alignments. For SINaTRA, we rank pairs by the reciprocal of the Euclidean distance between rank-normalized connectivity profiles. For BLAST, we rank pairs by bit-scores between protein sequences. We compute enrichment using area under the receiver operating characteristic curve (AUROC), average precision or area under the precision-recall curve (AUPR), and maximum *F*_1_ score (the harmonic mean of precision and recall) to take into account trade-offs between sensitivity and specificity. We note that precision-recall curves (summarized by the AUPR and maximum *F*_1_ score) have been cited to be more informative than ROC curves when the ratio of positives to negatives in a classification dataset is highly skewed or imbalanced [15]. Our classification dataset does have a large class imbalance with only 2.1% out of 35,846,366 human-mouse protein pairs classified as *k*-functionally similar at *k* = 100.

Table 1 shows the enrichment statistics for human and mouse proteins. We find that Handl better captures functional similarity than SINaTRA and HubAlign, and performs comparably to IsoRank and BLAST. Handl outperforms three of the four other benchmarked algorithms in AUPR by at least 10% (0.042 for Handl versus 0.038, 0.035, and 0.024 for IsoRank, HubAlign, and SINaTRA, respectively), and outperforms all other algorithms by at least 7% in maximum *F*_1_ score (0.089 for Handl versus 0.083, 0.081, 0.074, and 0.046 for BLAST, IsoRank, HubAlign, and SINaTRA, respectively). Interestingly, BLAST achieves the highest AUPR at 0.053. In terms of AUROC, Handl achieves an AUROC of 0.632 which is within 1% of the AUROC achieved by IsoRank (0.638), and is higher than HubAlign by 3% (0.611), BLAST by 12.5% (0.560), and SINaTRA by 23% (0.511). Thus, Handl compares favorably to standard approaches computing for cross-species gene/protein similarity.

**Table 1:** Results for *k*-functional similarity prediction between human-mouse using Handl and other algorithms. Notation for Handl indicate embeddings from *source → target*, i.e., 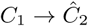 in Equation (1).

| Algorithm | AUROC | AUPR | Max $F_1$ |
| --- | --- | --- | --- |
| HANDL (human→mouse) | 0.630 | 0.042 | 0.088 |
| HANDL (mouse→human) | 0.632 | 0.042 | **0.089** |
| BLAST | 0.560 | **0.053** | 0.083 |
| ISORANK | **0.638** | 0.038 | 0.081 |
| HUBALIGN | 0.611 | 0.035 | 0.074 |
| SINATRA | 0.511 | 0.024 | 0.046 |

#### 2.4 Using Handl-homologs to find known and novel phenologs

In this section we show that Handl can yield new methods for *phenolog* discovery. The notion of a phenolog was defined in [38] as two phenotypes in different species that are related by the homology of their associated genes. The method used in [38] to identify phenologs is based on detecting an over-representation of homologous proteins associated with each phenotype in the pair. Reliance on homologs is a good start for determining phenotypic preservation, but the requirement of sequence preservation may be too restrictive if the primary goal is to study function across species [2,55,52].

Our goal is to investigate whether additional phenologs may be discovered through the expanded notion of homology that is provided by Handl (rather than reproduce the results in [38] with a different methodology). As a proof of concept, we show results using human-mouse Handl homology scores. To make consistent comparisons with [38] we use the same phenotype to gene association datasets as in that study.

To match the methods used in [38], we threshold Handl homology scores, leading to a binary classification of each cross-species gene pair as either a Handl-homolog or not. We set the threshold so as to output a small set consisting of the most confident phenolog predictions. To compare a phenotype in one species to a phenotype in a different species we count the number of Handl-homolog pairs across species in the two associated phenotypes. We then use the same procedure as in [38], with Handl-homologs playing the role that homologs did in that work; details are provided in Supplemental Information S1.3.

By using a stringent threshold on Handl homology scores, we sharply limit the size of the set of phenologs predicted. Whereas [38] reported 3634 phenotype pairs passing significance testing, our results show 47 pairs of phenotypes to be significantly associated. Within this set were 18 phenologs previously reported by [38], which is not surprising given that many homolog pairs are ranked highly by Handl (e.g. see Figure 2b). However our primary interest are the matches that are not part of the homolog-based phenologs reported in [38].

We find that Handl-based similarity can uncover many new phenologs that are not statistically significant when using homologs. As an example, we show in Figure 3 details of a phenolog found using Handl, but not found in [38]. This phenolog matches *Abnormal Muscle Fiber Morphology* in the mouse with *Muscular Dystrophy* in human. The example is illustrative as it shows that false negatives can occur when using only homoologs, even for straightforward matches such as this one. In fact the homolog-based method used in [38] finds only 1 homolog in common, while Handl detects 21 functionally-close pairs of proteins. The larger set of functionally-close protein pairs uncovered by Handl gives greater statistical power for cases such as this one, where there is only a small number of homologs shared between the two phenotypes.

**Fig. 3:**
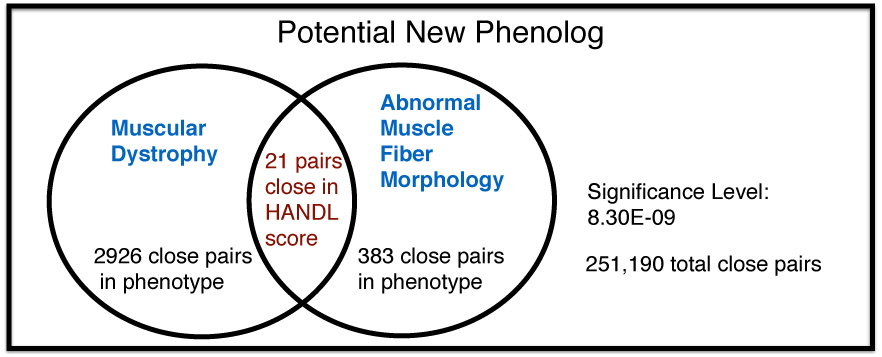
A potential new phenolog, not found by [38], relating the phenotypes Muscular Dystrophy (human) and Abnormal Muscle Fiber Morphology (mouse).

We find that many of the remaining 29 phenotype pairs identified using Handl are potentially valid phenologs. To demonstrate, we compare the human phenotype (disease) with mouse phenotype (symptom). The 29 pairs of phenotypes ranged over 8 unique human diseases and 24 unique mouse phenotypes. We obtained the description of each disease from the Genetics Home Reference^2^ and compared the disease description to the matched mouse phenotypes.

Table 2 groups the results into three categories: obvious symptom match, possible symptom match, and novel match. For each disease term and mouse phenotype pair, if the name of the mouse phenotype was indicated as a symptom in the disease description, we considered an obvious match; if the disease description contained a symptom that was similar to the mouse phenotype, we considered it a possible match; and if there was no similarity in the phenotype and the disease description we considered it as a novel match. Table 2 shows that, for over 75% of the new phenologs predicted using Handl, there is evidence in the literature supporting the association.

**Table 2:** Matched mouse phenotypes per human disease among newly-identified phenologs.

| Human Disease | Obvious Match | Similar Match | Novel Match |
| --- | --- | --- | --- |
| Systemic lupus erythematosus | 1 | 0 | 0 |
| Dilated cardiomyopathy | 4 | 0 | 1 |
| Zellweger syndrome | 2 | 1 | 2 |
| Dysfibrinogenemia | 0 | 0 | 1 |
| Myopathy | 1 | 0 | 0 |
| Bardet-Biedel syndrome | 6 | 2 | 1 |
| Adrenoleukodystrophy | 0 | 3 | 1 |
| Muscular Dystrophy | 2 | 0 | 1 |
| **Total** | **16** | **6** | **7** |

In summary, we find that Handl-homologous pairs can indeed provide a basis for expanding the set of phenologs discovered by previous methods. We anticipate future research developing methods to uncover phenologs may use previous methods in tandem with Handl-based methods to more thoroughly explore the space of possible phenologs.

### 2.5 Gene pairs with synthetic lethal interactions are co-located in Handl-space within and across species

In this section we show that Handl homology scores for gene pairs in different species are associated with pairwise gene function. We hypothesize that gene pairs with genetic interactions are separated in Handl-space from those without genetic interactions, and that the direction of separation is preserved when projecting other species into the same Handl-space. That is, gene pairs with genetic interactions in *P*_*C1*_ are co-located with gene pairs with genetic interactions in 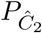 (see Section 4.3 for additional details). To test this hypothesis, we consider a particular form of genetic interaction – the synthetic lethal (SL) interaction – among genes in two different species of yeast: *S. cerevisiae* (*S.c.*) and *S. pombe* (*S.p.*).

We test whether SLs are separated from non-SLs in Handl-space, and whether this separation extends across species, by training a classifier for gene pairs using Handl-embeddings as features. More specifically, we train a random forest (RF) to classify gene pairs as SLs or non-SLs within both species *simultaneously*, using the source embedding (given by *P*_*C1*_) and the target embedded into source space (given by 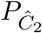). Without the Handl-embeddings, we would not be able to train a classifier for multiple species, since genes in different species would be in different vector spaces and have different dimensions. We perform 4-fold cross-validation, fixing the relative fraction of pairs from each species, and assess the degree of separation between SLs and non-SLs in Handl-space by evaluating the RF classifications with maximum *F*_1_ score (the harmonic mean of precision and recall), the area under the ROC curve (AUROC), and the area under the precision-recall curve (AUPRC). We report the average across the four folds, separating the results by species. We use a nested cross-validation strategy to choose the number of trees for the RF that maximizes the held-out AUPRC. For simplicity, all of our experiments in this section use *S.c.* as the source and *S.p.* as the target.

We first train classifiers on an SL dataset of high-confidence, low-throughput interactions from BioGRID [9] (see Section 4.4 for additional details of the dataset). This dataset is the most recent update of the dataset used by Jacunski, et al. [26], and we follow their approach by creating a dataset with an equal number of non-SLs sampled randomly from pairs in the PPI networks without SLs. Table 3 shows the results, which demonstrate that SLs are significantly separated from non-SLs in Handl-space. The RFs achieve average AUROC of 0.933 in *S.c.* (0.876 in *S.p.*), AUPRC of 0.933 (0.877), and maximum *F*_1_ score of 0.860 (0.814). We then compare the results using Handl features to SINaTRA (which also uses a RF), using the same protocol as above but train RFs with the rank-normalized features produced by SINaTRA for cross-species predictions in [26]. SINaTRA achieves average AUROC of 0.908 in *S.c.* (0.880 in *S.p.*), AUPRC of 0.907 (0.892), and maximum *F*_1_ score of 0.834 (0.808). Thus, while both sets of classifiers make highly accurate predictions, the RFs trained using Handl features outperform SINaTRA on four of six measures.

**Table 3:** Results training classifiers for synthetic lethal interactions on baker’s yeast (*S.c.*) and fission yeast (*S.p.*) data *simultaneously*. We compute performance separately for each species (indicated by “Test species”). For each statistic, we report the average on held-out data from 4-fold cross-validation, and bold the highest (best) score.

| Dataset | Test species | Features | AUROC | AUPRC | Max $F_1$ |
| --- | --- | --- | --- | --- | --- |
| BioGRID [9] | *S.c.* | HANDL | **0.933** | **0.933** | **0.860** |
|  |  | SINATRA | 0.908 | 0.907 | 0.834 |
|  | *S.p.* | HANDL | 0.876 | 0.877 | **0.814** |
|  |  | SINATRA | **0.880** | **0.892** | 0.808 |
| Chromosome biology [13,56] | *S.c.* | HANDL | **0.864** | **0.402** | **0.421** |
|  |  | SINATRA | 0.788 | 0.201 | 0.282 |
|  | *S.p.* | HANDL | 0.822 | 0.370 | 0.423 |
|  |  | SINaTRA | **0.837** | **0.393** | **0.437** |

Next, we train linear support vector machines (SVMs) instead of RFs to learn hyperplanes separating SLs and non-SLs in Handl-space, and only see a small degradation in performance compared to the RF (Table S4). This further supports the case that SLs in different species are co-located in Handl-space because, unlike the decision boundary learned by the RF, the SVM learns a linear decision boundary. Therefore, because the linear SVM classifies SLs with high accuracy in both species simultaneously, there is evidence for a direction in the Handl-embeddings capturing synthetic lethality.

We then further evaluate the separation of SLs and non-SLs in Handl-space and train random forest classifiers with matched high-throughput datasets from *S.c.* and *S.p.* These datasets consist of SL and non-SL pairs among 743 *S.c.* genes [13] and 550 *S.p.* genes [56] involved in chromosome biology (see Section 4.4 for additional details of the dataset). The key differences between the chromosome biology SL datasets and the BioGRID datasets are that the chromosome biology datasets are restricted to functionally similar genes, include 5.5% SLs and 94.5% measured non-SLs in *S.c.* and 10.6% SLs and 89.4% measured non-SLs in *S.p.* (unlike the BioGRID data which only measured SLs), and were generated through high-throughput experiments.

Table 3 shows that the RFs trained on Handl-embeddings achieve significant predictive performance on held-out data from the chromosome biology dataset, with an AUROC of 0.864 in *S.c.* (0.822 in *S.p.*), AUPRC of 0.402 (0.370), and maximum *F*_1_ score of 0.421 (0.423). While these results show significant predictive power and also indicate that SLs are co-located across species in Handl-space, the performance of the classifiers is much poorer than on the BioGRID data. This is likely due to a combination of the noisy, high-throughput nature of the data measurements and the class imbalance. Interestingly, on the chromosome biology dataset, Handl outperforms SINaTRA by a large margin for predictions in *S.c.*, while SINaTRA outperforms Handl by a smaller margin for predictions in *S.p.*

The significant precision and recall these classifiers achieve using Handl-embeddings stand in stark contrast to the results achieved using sequence homologs to transfer knowledge. Following earlier approaches [16], we predicted SLs across species using orthologs, obtaining a single point on the ROC (or precision-recall) curve. Specifically, for a pair (*u, v*) in the target species, we predicted (*u, v*) to have a SL if *u* and *v* each have orthologs *u^′^* and *v^′^* in the source species, respectively, and (*u^′^, v^′^*) is SL in the source species. On the BioGRID dataset, this approach achieves precision of 1.0 but recall of only 0.005 resulting in a maximum *F*_1_ score of 0.01, and on the chromosome biology dataset, this approach achieves modest precision of 0.269, but recall of only 0.002, resulting in a maximum *F*_1_ score of 0.005. Because of the limited number of homolog pairs, this approach predicts many false negatives and provides no insight of epistatic relationships between genes without homologs.

Together, these results not only show that synthetic lethal interactions are significantly clustered *across species* in the Handl-embeddings but also show that Handl-embeddings can leverage knowledge of homologous genes across species to transfer knowledge and gain insight on other genes within species.

### Evaluating the effect of pair-inputs on the classifiers

Predicting synthetic lethal interactions between gene pairs using features constructed for individual genes is an example of a pair-input classification problem. A challenge with evaluating classifiers trained on pair inputs with held-out data is that, for a given pair (*u, v*), it is possible that the features for only *u*, only *v*, both *u* and *v*, or neither *u* and *v*, can be found in the training data [49]. Thus, information concerning genes found in the held-out data may be “leaked” to the classifier during training. To evaluate the effect of this issue, Park & Marcotte [49] suggest evaluating classifications for gene pairs which contain one, two, or no genes in the training data separately. This is analogous to holding out individual *genes* instead of *gene pairs* at training time and, thus, we evaluate the effect of pair-inputs by repeating the experiments above but hold out genes instead of gene pairs for evaluation. We report the results in Table S5. We find that the classifiers are able to predict SLs for genes *not* found in the training data, but with a significant change in performance. On the BioGRID dataset, the classifiers achieve an AUROC of 0.872 in *S.c.* (0.823 in *S.p.*), an AUPRC of 0.875 (0.814), and maximum *F*_1_ of 0.797 (0.772). On the chromosome biology dataset, the classifiers achieve an AUROC of 0.701 in *S.c.* (0.691 in *S.p.*), an AUPRC of 0.160 (0.207), and maximum *F*_1_ of 0.202 (0.285). We hypothesize that the larger drop in performance on the chromosome biology data is due to the matched nature of the *S.c.* and *S.p.* datasets. We also find similar drops in performance for SINaTRA when holding out genes instead of pairs (also in Table S5).

### 2.6 Leveraging multiple model organisms for function prediction

As a final exploration of the utility of Handl for cross-species functional inference, we study the potential for leveraging annotations from multiple model organisms simultaneously. This problem has not previously been explored extensively, with the exception of [42], which took a Bayesian approach. Our intent is not to propose a new method for function prediction, but to demonstrate the value of cross-species information as obtained via Handl.

We focus on function prediction in three species: human, mouse, and baker’s yeast (*S.c.*). Following the approach taken in [35], we focus on predicting rare GO terms – in particular we form predictions for all GO terms that occur between 2 and 300 times in the annotation corpus of one species. We study the information content of multiple Handl scores by constructing a convex combination *α_H_, α_Y_, α_M_* of a given species’ diffusion score (*D*_1_) with the Handl homology scores (*D*_12_) of the two other species.

We perform a binary classification for each GO term. We rank proteins by the total scores contributed by other proteins – within the same and across species – annotated with that term, weighted by *α_H_, α_Y_, α_M_*. We then assess performance using the maximum *F*_1_ score averaged over all GO terms. We test whether such a convex combination has greater predictive power for functional inference than just using information from any single species. Details of our method are given in Supplemental Information S1.2.

Figure 4 shows the prediction performance obtained over the simplex (*α_H_, α_Y_, α_M_*) such that ∑ *α_i_* = 1. Contours show *F*_1_ scores, and the point of maximum *F*_1_ score is plotted. For comparison purposes we provide results for the case in which Handl scores are randomly permuted. The figure shows that in each case, the greatest improvement in functional prediction comes when making use of information from both additional species. The improvements in functional prediction are shown in Table S6.

**Fig. 4:**
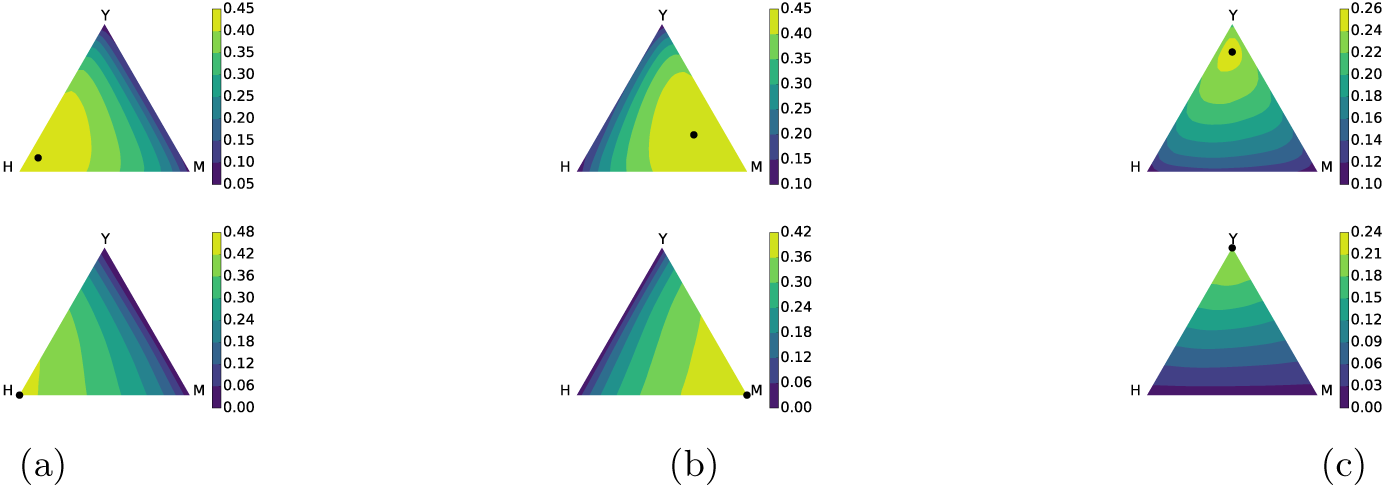
*F*_1_ score of functional inference using multiple additional species. Inferring protein functions for (a) human – H, (b) mouse – M, and (c) baker’s yeast – Y. Upper row: using Handl scores; Lower row: null hypothesis, using randomly permuted Handl scores.

## 3 Discussion

We introduce a novel, kernel-based algorithm to embed proteins from different species into a shared vector space. Our approach combines standard sequence homology with the idea of network homology through the use of network *landmarks*. We use a particular kernel – the regularized Laplacian – to create a functional embedding, and use the resulting algorithm, Handl, to embed proteins from humans, mice, and yeast into shared spaces. We interpret inner products in this vector space as homology scores, and show that the embedding itself is biologically meaningful.

We evaluate Handl embeddings through comparisons with other types of genetic and network homology on a variety of cross-species tasks, validating predictions from Handl with the biological literature and held-out data. First, we show that Handl homology scores are strongly correlated with functional similarity. We then use Handl embeddings for two cross-species tasks traditionally performed using homologs: identifying “orthologous phenotypes” (phenologs) [38] and classifying synthetic lethal interactions. The phenologs identified using Handl include 18 previously discovered [38], but importantly include 29 new phenologs that were not statistically significant using orthologs. Most of these new phenologs missed by earlier methods are supported by the biological literature. We also find Handl encodes pairwise gene function, as pairs with synthetic lethal interactions are co-located in Handl-space, both within and across species. We also demonstrate how function prediction methods that integrate data from multiple species may outperform methods that only use data from a single species. Importantly, in these tasks, we transfer knowledge both from humans to model organisms – as is commonly done to derive disease models – and from model organisms to humans – as is commonly done for comparative genomics. Thus, Handl represents a new direction towards realizing the crucial goal of algorithms for transferring knowledge of genetics across species.

We anticipate that combining network landmarks and diffusion kernels could also be useful for a broader class of data-driven tasks. The kernel used by the Handl algorithm in this study is based on the “guilt-by-association” hypothesis for functional annotation, but it may be possible to learn how to use this information for different prediction tasks. For example, many recent methods have successfully used vectors of diffusion-based similarity scores as features for off-the-shelf supervised learning [8,11], and achieved state-of-the-art performance on multiple tasks [11]. Seen in that light, the Handl embeddings can be seen as a component of a *transfer learning* [48] approach for cross-species inference. Transfer learning – using knowledge gained in solving one task to aid in solving a different task – is often approached by finding appropriate transformations of data features. These transform a source dataset in a fashion such that a model trained on the source data can be usefully applied to a different, target dataset (‘domain adaptation’, e.g., see [61]). Methods for aligning the source and target embeddings may be required to make such a transfer learning approach possible. At the same time, we showed that methods for transferring knowledge across species can be useful even when there is a wealth of data in the target species. Thus, to achieve optimal performance, supervised learners may need to train on multiple species *simultaneously*.

Beyond kernels derived from protein interaction, there are a wide range of other kernels that can inform biological function assessment, including kernels derived from co-expression, genetic interaction, metabolic pathways, domain structure, and sequence [33,8,34]. Because Handl is a method for creating a new kernel encompassing the nodes of multiple networks, it holds potential as a new tool for kernel learning methods such as support vector machines in a wide variety of applications beyond cross-species function prediction.

## 4 Methods

### 4.1 Multi-Species Network Kernel Embedding

As described in Section 2, Handl constructs a multi-species network kernel embedding. Here we provide additional details on the method.

We start by noting that there are a large variety of kernels derived from networks [31], [19, Ch. 2], and they can model processes such as random walks, heat diffusion, PageRank, electrical resistance, and other ways of capturing node similarity in a network. Many kernels derived from networks have been applied successfully for a wide range of problems associated with biological network analysis (e.g., see review in [14]).

Though many previous studies have used graph kernels to compare nodes *within* biological networks, to our knowledge, few methods have utilized kernels to fulfill the goal of comparing nodes *across* multiple biological networks. This is the challenge that the Handl algorithm addresses.

To do so, Handl relies on a basic property of a kernel: any kernel is also an *inner product* in a particular space. That is, for any kernel *κ*(*⋅, ⋅*), there is a function *ϕ*(*⋅*) that assigns vectors to nodes such that that *κ*(*i, j*) = *ϕ*(*i*)^T^ *ϕ*(*j*). The corresponding vector space (termed the *reproducing kernel Hilbert space* (RKHS)) introduces a geometric interpretation for the kernel function. In the context of a kernel for network nodes, the RKHS representation can be thought of as an *embedding* of the network into a vector space in a manner that captures node similarity via inner product.

Conceptually, Handl approaches the multi-network challenge by constructing a *joint embedding* of the nodes of two (or more) networks in a *single* RKHS. The key to the Handl method is that, within this RKHS, the similarity of nodes from *different* networks is still captured by inner product, resulting in a *multi-network kernel*.

Given a *source* network *G*_1_, a *target* network *G*_2_, and a kernel *κ*, the strategy taken by Handl is to start by embedding the nodes of the source network *G*_1_ using the associated function, *ϕ*. As described above, this means that inner product between embedded *G*_1_ nodes will capture similarity as described by *κ*. Next, Handl makes use of *landmarks* – pairs of nodes in the source and target networks with identical function. The idea is to embed the nodes of the target network *G*_2_ into the same space as the nodes of the source network *G*_1_, such that their position in that space reproduces their similarity to the landmarks of *G*_2_. Essentially, we posit that locating a node from *G*_2_ based on its similarity to the set of landmarks in *G*_1_ will also establish its similarity with the *non-landmark* nodes in *G*_1_. As a result, Handl creates a multi-network kernel – a single kernel function that captures both the similarity of nodes to each other in the source network *G*_1_, and the similarity of nodes between the source and target networks *G*_1_ and *G*_2_.^3^

We now define the Handl approach formally. Let the matrix *K ∈* ℝ^*n×n*^ hold the values of the similarity function *κ*(*i, j*) for all pairs of *n* proteins from a particular species. For any such kernel matrix, we can write *K* = *CC^T^* where *C* is an *n × k* matrix, uniquely defined up to an orthogonal transformation, with *k ≤ n*. This follows from the fact that *K* is positive semidefinite, and means that 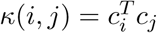, where *c_i_* is the *i*th row of *C*, represented as a column vector. As explained above, the similarity between nodes *v*_*i*_ and *v_j_* is exactly given by the inner product of their corresponding vectors, *c_i_* and *c_j_*.

Conceptually, Handl approaches the multi-network challenge by constructing a *joint embedding* of the nodes of two (or more) networks in a *single* RKHS. The key to the Handl method is that, within this RKHS, the similarity of nodes from *different* networks is still captured by inner product, resulting in a *multi-network kernel*.

Now consider a source network *G*_1_ = (*V*_1_*, E*_1_) and a target network *G*_2_ = (*V*_2_*, E*_2_) with *|V*_1_*|* = *m* and *|V*_2_*|* = *n*. We assume the existence of some (small set of) nodes that correspond between *G*_1_ and *G*_2_. In the case where *G*_1_ and *G*_2_ are PPI networks, these can be orthologous proteins. For example, for orthologous proteins in different networks, it is well known that evolutionary rates differ over a wide range of magnitudes [7]. Some proteins are highly conserved, and their orthologs will have substantial sequence similarity between *G*_1_ and *G*_2_. Thus, there are generally a small subset of proteins that can be confidently mapped between *G*_1_ and *G*_2_ based on the magnitude and uniqueness of the similarity of their sequence information which we refer to as landmarks.

We then proceed as follows. First, we construct kernel (similarity) matrices *D*_1_ *∈* ℝ^*m×m*^ and *D*_2_ℝ^*n×n*^ corresponding to *G*_1_ and *G*_2_. Next, we construct RKHS vector representations *C*_1_ for nodes in the source network *G*_1_ from the factorization 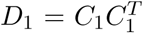. Let *C*_1*L*_be the subset of the rows of *C*_1_ corresponding to landmarks, and let *D*_*2L*_ be the subset of the rows of *D*_2_ corresponding to landmarks (in corresponding order).

The key step then is to construct the vector representations of the nodes in the target network *G*_2_. To do this, we treat the similarity scores *D*_*2L*_ in the target network as if they applied to the landmarks in the source network *G*_1_. For a given node in the target network, we want to find a vector for the node such that its inner product with each *source* landmark vector is equal to its diffusion score to the corresponding *target* landmark. This implies that the RKHS vectors, 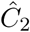, for nodes in the target network *G*_2_ should satisfy 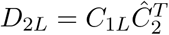. This underdetermined linear system has solution set,

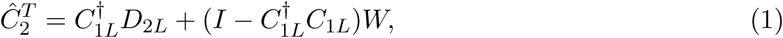

where 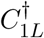 is the Moore-Penrose pseudoinverse of *C*_*1L*_, and *W* is an arbitrary matrix. We choose the solution corresponding to *W* = 0, meaning that the vectors 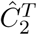 are the solutions having minimum norm.

The resulting solution, 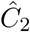, represents the embedding of the nodes of *G*_2_ (the target) into the same space as the nodes of *G*_1_ (the source). We can then compute similarity scores for all pairs of nodes across the two networks as 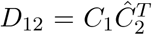. This yields *D*_12_, an *m × n* matrix of similarity scores between nodes in the source and target networks.

### 4.2 Handl and the Regularized Laplacian

While our method can be used with any kernel, in this paper we focus on using a kernel intended to capture functional similarity of proteins. To motivate our choice of kernel function *k*(*⋅, ⋅*), we consider the function prediction problem on a single network, *G* = (*V, E*), with *|V |* = *n*, where *G* has adjacency matrix *A* with entries *a_ij_*. For simplicity we consider *G* to be unweighted, so *a*_*ij*_ ∈ {0, 1}. Extensions of our arguments to weighted graphs are straightforward.

A central idea used throughout network-based functional prediction methods is that of *guilt by association* – that is, two nodes that are near each other are more likely to share the same label than two nodes that are far apart. In the context of protein function prediction, this principle has been well established. For example, the authors in [35] show that two neighbors in the protein interaction network are more likely to have the same function than a randomly chosen pair.

Consider the case of determining whether nodes should receive a particular function label where we label a node with a 1 if it should receive the label, and 0 otherwise. We are interested in the case in which the label is rare, and we believe that nodes may be mislabeled (e.g., some nodes labeled 0 should actually be labeled 1). We assume that there is some current labeling which is incomplete; that is, most nodes are currently labeled 0, and some nodes labeled 0 should actually be labeled 1. Define the vector *y* such that *y_i_* = 1 if node *v*_*i*_ has label 1, and zero otherwise. The goal of the function prediction problem is to estimate a new 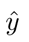 that is a better labeling of nodes in *V*.

To address this problem, we proceed as follows [5,18]. First, we posit that *y* should not differ too much from 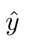 nodes should tend to be given the labels they already have. Second, we also posit that neighbors in *G* should tend to be given the same label – this is the guilt by association principle.

Note that these two goals are in conflict: fully following the first principle leaves all labels unchanged, while fully following the second principle makes all labels the same (either 0 or 1). To balance these, we define the following optimization:

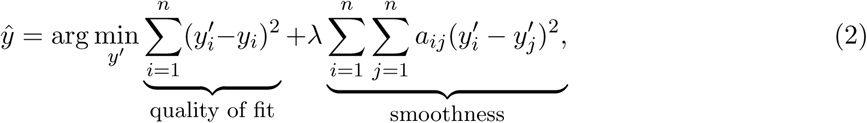

in which we use *λ* to control the tradeoff between the two principles.

This expression can be compactly expressed using the Laplacian of *G*: *L* = *D − A* in which *D* is a diagonal matrix with node degree on the diagonal: *D_ii_* = ∑_*j*_*a*_*ij*_. Then,

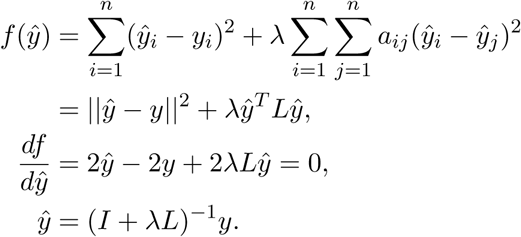

The matrix (*I* + *λL*)^−1^ is the *regularized Laplacian* of *G* [67]. It is a positive semidefinite matrix and hence a kernel.^4^

The combination of the multi-network kernel embedding (Section 4.1) with the Regularized Laplacian constitutes Handl, and (as noted above) the resulting cross-species similarity scores are Handl *homology scores.* We denote the Handl homology score of two proteins *p*_*i*_ and *p*_*j*_ as *d_ij_*, we refer to the RKHS in which *G*_1_ and *G*_2_ are embedded as Handl*-space*, and the Handl-embeddings are given by the rows of *C*_1_ and 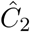 (Section 4.1).

### 4.3 Scores for gene pairs

We also find it useful to develop a measure to capture functional similarity between two *pairs* of nodes across species. Pair-similarity can then be used to predict outcomes for pairs of genes (e.g. synthetic lethality). Given two pairs of nodes (*v_i_, v_j_*), (*v_k_, v_ℓ_*), we define a pairwise similarity metric such that the score is large only if *d_ik_* and *d_jℓ_* (or *d_iℓ_* and *d_jk_*) are both large. This reflects the hypothesis that synthetic lethal interactions occur within pathways, and between pathways that perform the same/similar essential biological function [6,40].

Hence, to compare pairs, we simply sum the Handl-embeddings for the nodes in each pair, and then compute Handl scores in the usual way. Given a matrix *C* of Handl-embeddings for nodes, we define the Handl-embeddings for a *pair* of nodes (*v_i_, v_j_*) as *P_C_*(*v_i_, v_j_*) = *c_i_* + *c_j_*. Computing similarity for a two pairs (*v_i_, v_j_*) and (*v_k_, v_ℓ_*) then yields:

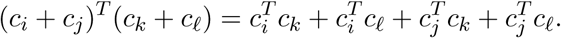

In general we expect each of the terms on the right hand side to be close to zero *unless* there is functional similarity between the corresponding nodes, because in high dimension, independent random vectors tend to be nearly orthogonal. We note that the pair-similarity scores and the pair-embeddings themselves can be used for a variety of tasks.

The utility of this approach is informed by recent work on predicting synthetic lethal interactions from network diffusion [11].

### 4.4 Data

#### Sequence homologs

For each combination of two organisms, we identify homologous pairs of genes using NCBI’s Homologene database [57] and chose 400 homolog-pairs at random, which constitute our landmarks.

#### Protein-protein interaction networks

We constructed protein-protein interaction (PPI) networks in *S.c.*, *S.p.*, mouse, and human (refer to Supplemental Information S2.1 for details). We restricted each network to the two-core of the largest connected component, and report summary statistics of the networks in Table S1. We use the two-core of the graph because topologically indistinguishable nodes (nodes that participate in an automorphism of the graph) will necessarily have identical Handl homology scores. A large class of topologically indistinguishable nodes includes many of the leaf nodes in the graph (degree-1 nodes). That is, in any case where there are two or more leaf nodes attached to the same parent, the nodes are topologically indistinguishable. Protein names were standardized by mapping to UniProt Accession IDs using mappings provided by the UniProt consortium [63].

#### Protein function annotations

Protein functions were determined using the Gene Ontology database (GO) [4]. Currently, GO contains more than 40,000 biological concepts over three domains: Molecular Function (MF), Biological Process (BP) and Cellular Component (CC). We use GO annotation corpora downloaded from SGD [10] for yeast, and UniProt [63] for all other species. We exclude annotations based on IEA or IPI evidence due to their lower associated confidence levels.

#### Synthetic lethal interactions

We constructed datasets of synthetic lethal (SL) interactions and non-interactions (non-SLs) from two high-throughput studies of analogous proteins in baker’s (*S.c.*) and fission (*S.p.*) yeast [13,56]. Also, following Jacunski, et al. [26], we constructed datasets of SL interactions from BioGRID (v3.4.157) in *S.c.* and *S.p.*, sampling an equivalent number of non-SLs from pairs in the PPI network without an SL. We report the size of the datasets in Table S2, and additional details in the Supplemental Information S2.2.

## Acknowledgements

This work was supported in part by NSF grants IIS-1421759 and CNS-1618207 (to M.C.) and by a grant from the Boston University Undergraduate Research Opportunities Program (to T.L.). We thank Simon Kasif, Evimaria Terzi, Prakash Ishwar, Lenore Cowen, Donna Slonim, and the Tufts BCB group for helpful discussion on this work.

## Supplemental Information

### S1 Methods

#### S1.1 Parameter choice

For all regularized Laplacians, we used a value of *λ* = 0.05. We found that the resulting relationship between Resnik similarity and Handl homology score did not vary significantly for *λ* values between 0.005 and 0.1.

#### S1.2 Function Prediction Methods

We assess prediction accuracy using leave-one-out cross validation. Let 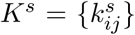 denote the regularized Laplacian for species *s*. For GO term *g*, let 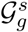 be the set of proteins in species *s* that are annotated with *g*. The same-species annotation score for a given protein *p* and GO term *g* is

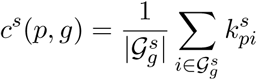

in which *p* is excluded from the sum (i.e., if it is contained in 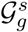). We also construct a cross-species annotation score for each protein, in which Handl scores with respect to proteins in the other organism are used:

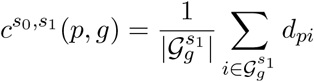

where *d_pi_* = *D*_12_(*p, i*) is the Handl score for protein *p* in species *s*_0_ and protein *i* in species *s*_1_. The prediction score is then *h*(*p, g*) = *α c^s^*^0^ (*p, g*) + (1 *− α*)*c^s^*^0*,s*1^ (*p, g*). To use multiple cross-species annotations, say *n*, we generalize *h*(*p, g*) to a convex combination of the same- and cross-species annotation scores: *h*(*p, g*) = *α*_0_ *c^s^*^0^ (*p, g*) + 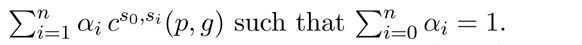.

We evaluate predictions using area under the receiver operating curve (AUC) and maximal F-score (over all detection thresholds). Since we are concerned with predicting rare GO terms, we find that maximal F-score is generally a more discriminative metric. We set the convex coefficients *{α_i_}* via cross-validation.

#### S1.3 Phenolog Discovery

Our method matches that used in [38], using protein pairs with high Handlhomology scores rather than homologs obtained from Homologene.^5^ Specifically, let *P*_1_ be the genes associated with the phenotype in species 1 and *P*_2_ be the genes associated with the phenotype in species 2. Our contingency table consists of the counts of the number of Handl-homologs involving *P*_1_ *∩ P*_2_, *P*_1_ *\ P*_2_, *P*_2_ *\ P*_1_, and (*Ω \ P*_1_) *\ P*_2_, with *Ω* denoting the set of all close pairs. We used a Fisher exact test to measure significance, and considered the match significant if the uncorrected *P*-value was less than 0.05. We corrected for multiple testing using a Bonferroni correction; there were 1, 278, 312 possible phenotype matches so we set the significance level at 3.9 × 10^−8^.

### S2 Data

#### S2.1 Protein-protein interaction networks

We constructed protein-protein interaction (PPI) networks in *S.c.*, *S.p.*, mouse, and human. The *S.c.* and *S.p.* networks were obtained from the Biological General Repository for Interaction Datasets (BioGRID) [9] version 3.4.157. Mouse and human PPIs were obtained from the STRING database version 9.1 [20]. PPI networks obtained were processed by mapping the protein names to the same namespace. Genes that could not be mapped via the UniProt database were removed from the PPI networks entirely. We provide further details of the network processing in Section 4.4. Table S1 shows summary statistics for the PPI networks before and after processing.

#### S2.2 Synthetic lethal interactions

We constructed datasets of synthetic lethal (SL) interactions in *S.c.* and *S.p.* from published epistatic miniarray profiles (E-MAPs). E-MAPs include genetic interactions scores for pairs of genes, where the magnitude of the score reflects the strength of the genetic interaction. We downloaded E-MAPs for *S.c.* from the supplementary information of Collins, et al. [13], and for *S.p.* from the supplementary information of Roguev, et al. [56]. We classified each pair of genes in each E-MAP as SL, non-SL, or uncertain. We used the thresholds from the Collins, et al. [13] supplementary information to classify pairs in *S.c.*. Given a pair with E-MAP score *ϵ*, we classified it as SL if *ϵ < −*3, uncertain if −3 *≤ ϵ < −*1, and non-SL otherwise. Similarly, we used the threshold for synthetic lethality from the Roguev, et al [56] supplementary information and used the same threshold for uncertainty, classifying *S.p.* pairs as SL if *ϵ < −*2.5, uncertain if −2.5 *≤ ϵ < −*1, and non-SL otherwise. We also remove pairs of genes in which either gene is not found in the corresponding PPI networks described in Sec 4.4. The resulting datasets included 7,165 SL and 123,507 non-SL interactions in *S.c.*, and 5,599 SL and 97,541 non-SL interactions in *S.p*.

We also constructed datasets of SL interactions from BioGRID [9] (v3.4.157)^6^. For both *S.c.* and *S.p.*, we extracted interactions of type “Synthetic Lethality” only (i.e. ignoring other negative genetic interactions), yielding datasets of 13,645 and 908 SLs, respectively.

We then standardized the datasets by mapping genes names to Uniprot Accession IDs [63]. Genes that could not be mapped via UniProt were excluded for this study, as were those that were not found in the processed PPI networks. For the BioGRID SL dataset, we followed Jacunski, et al. [26], by sampling an equivalent number of non-SLs pairs from genes PPI networks that do not partake in SL interactions in the BioGRID SL dataset. Table S2 shows summary statistics of the SL datasets before and after processing.

### S3 Results

#### S3.1 Associations of Handl scores with functional similarity for other pairs of species

We associated Handl homology scores and functional similarity for pairs of proteins in additional pairs species, using the methodology described in Section 2.3. Figures S1 and S2 show the results for embedding human into yeast, and mouse into yeast.

#### Footnotes

1 A pair of genes is *homologous* if the pair share a common ancestor – the pair is called *orthologous* if they are in different species [17].

2 https://ghr.nlm.nih.gov/

3 We note that this is a fundamentally different strategy than has been used in past manifold-alignment methods [22,66], in which alignment is based on Euclidean distance. Aligning on Euclidean distance does not respect inner product, and so the similarity captured by the kernel is not preserved in the alignment.

4 In addition to the “guilt by association” argument, we note an additional reason from [64,8] that the regularized Laplacian is an appropriate tool for functional inference on protein interaction networks: it also naturally discounts paths that pass through high-degree nodes.

5 Note that none of the landmarks (which are a subset of the homologs) are used to discover new phenologs.

6 https://thebiogrid.org

## S4 Supplementary Tables

**Table S1:**
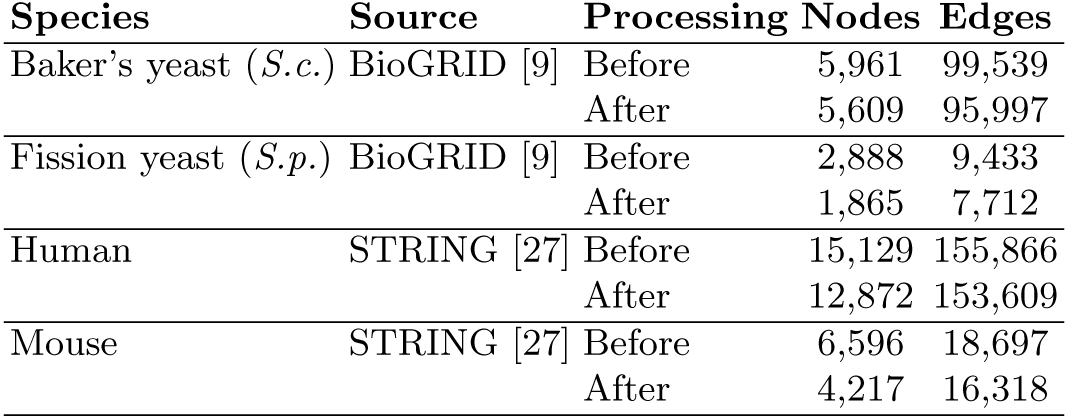
Summary statistics of PPI networks. We processed the graphs to restrict to the two-core of the largest connected component.

**Table S2:**
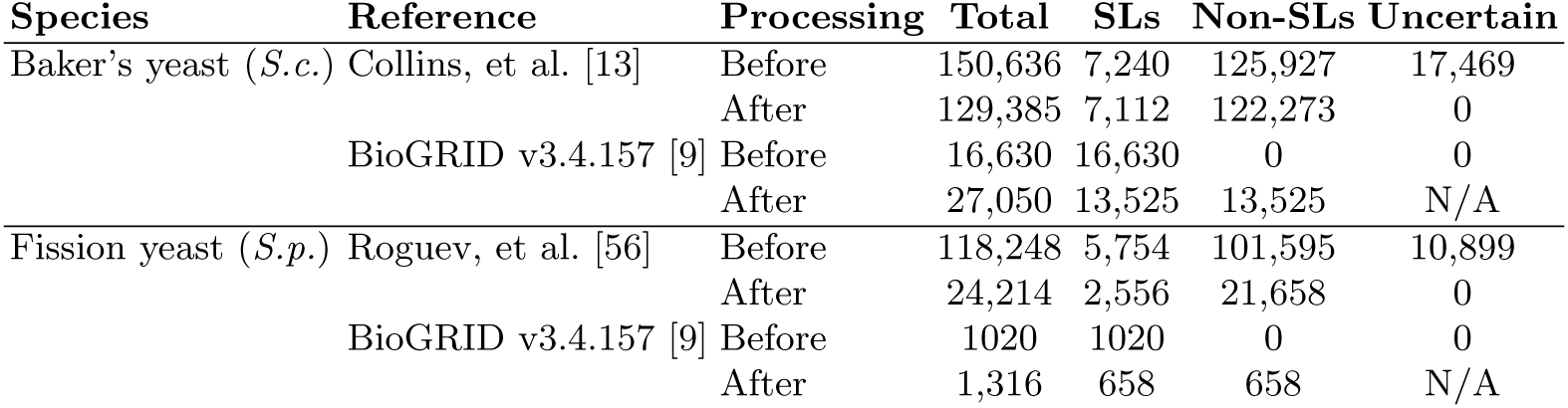
Summary statistics of synthetic lethal interaction datasets.

| Species | Reference | Processing | Total | SLs | Non-SLs | Uncertain |
| --- | --- | --- | --- | --- | --- | --- |
| Baker's yeast (*S.c.*) | Collins, et al. [13] | Before | 150,636 | 7,240 | 125,927 | 17,469 |
|  |  | After | 129,385 | 7,112 | 122,273 | 0 |
|  | BioGRID v3.4.157 [9] | Before | 16,630 | 16,630 | 0 | 0 |
|  |  | After | 27,050 | 13,525 | 13,525 | N/A |
| Fission yeast (*S.p.*) | Roguev, et al. [56] | Before | 118,248 | 5,754 | 101,595 | 10,899 |
|  |  | After | 24,214 | 2,556 | 21,658 | 0 |
|  | BioGRID v3.4.157 [9] | Before | 1020 | 1020 | 0 | 0 |
|  |  | After | 1,316 | 658 | 658 | N/A |

**Table S3:** Results for *k*-functional similarity prediction between *S.c.*-*S.p.* using Handl and other algorithms. Notation for Handl indicate embeddings from *source → target*, i.e., 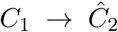 in Equation (1). Between *S.c.* and *S.p.*, only 1.18% of 6,483,610 possible protein pairs are labeled to be *k*-functionally similar at *k* = 100.

| Algorithm | AUROC | AUPR | Max $F_1$ |
| --- | --- | --- | --- |
| HANDL (*S.c.*→*S.p.*) | **0.633** | 0.026 | 0.061 |
| HANDL (*S.p.*→*S.c.*) | 0.630 | 0.032 | 0.068 |
| BLAST | 0.565 | **0.063** | **0.124** |
| ISO RANK | 0.607 | 0.018 | 0.040 |
| HUBALIGN | 0.551 | 0.015 | 0.036 |
| SINATRA | 0.524 | 0.013 | 0.026 |

**Table S4:** Results training linear support vector machines to classify synthetic lethal interactions on *S.c.* and *S.p.* data *simultaneously*. We compute performance separately for each species (indicated by “Test species”). For each statistic, we report the average on held-out data from 4-fold cross-validation over *gene pairs*, and bold the highest (best) score.

| Dataset | Test species | Algorithm | AUROC | AUPRC | Max $F_1$ |
| --- | --- | --- | --- | --- | --- |
| BioGRID [9] | *S.c.* | HANDL | **0.953** | **0.933** | **0.892** |
|  |  | SINATRA | 0.798 | 0.774 | 0.750 |
|  | *S.p.* | HANDL | 0.714 | 0.657 | 0.731 |
|  |  | SINATRA | **0.786** | **0.846** | **0.850** |
| Chromosome biology [13,56] | *S.c.* | HANDL | **0.865** | **0.334** | **0.396** |
|  |  | SINATRA | 0.631 | 0.092 | 0.156 |
|  | *S.p.* | HANDL | **0.711** | 0.123 | **0.208** |
|  |  | SINATRA | 0.569 | **0.124** | 0.214 |

**Table S5:** Results training random forests to classify synthetic lethal interactions on *S.c.* and *S.p.* data *simultaneously*. We compute performance separately for each species (indicated by “Test species”). For each statistic, we report the average on held-out data from 4-fold cross-validation over *genes*, and bold the highest (best) score.

| Dataset | Test species | Algorithm | AUROC | AUPRC | Max $F_1$ |
| --- | --- | --- | --- | --- | --- |
| BioGRID [9] | *S.c.* | HANDL | **0.872** | **0.875** | **0.797** |
|  |  | SINATRA | 0.848 | 0.852 | 0.779 |
|  | *S.p.* | HANDL | 0.823 | 0.814 | **0.772** |
|  |  | SINATRA | **0.839** | **0.855** | 0.769 |
| Chromosome biology [13,56] | *S.c.* | HANDL | **0.701** | **0.160** | **0.202** |
|  |  | SINATRA | 0.681 | 0.115 | 0.184 |
|  | *S.p.* | HANDL | 0.691 | 0.207 | 0.285 |
|  |  | SINATRA | **0.723** | **0.239** | **0.314** |

**Table S6:** Improvement in functional prediction using two other species.

| Target Species | AUC | $F_1$ score |
| --- | --- | --- |
| Human | 0.3% | 2.6% |
| Mouse | 1.0% | 8.6% |
| Baker’s yeast | 0.3% | 16.0% |

## S5 Supplementary Figures

**Fig. S1:**
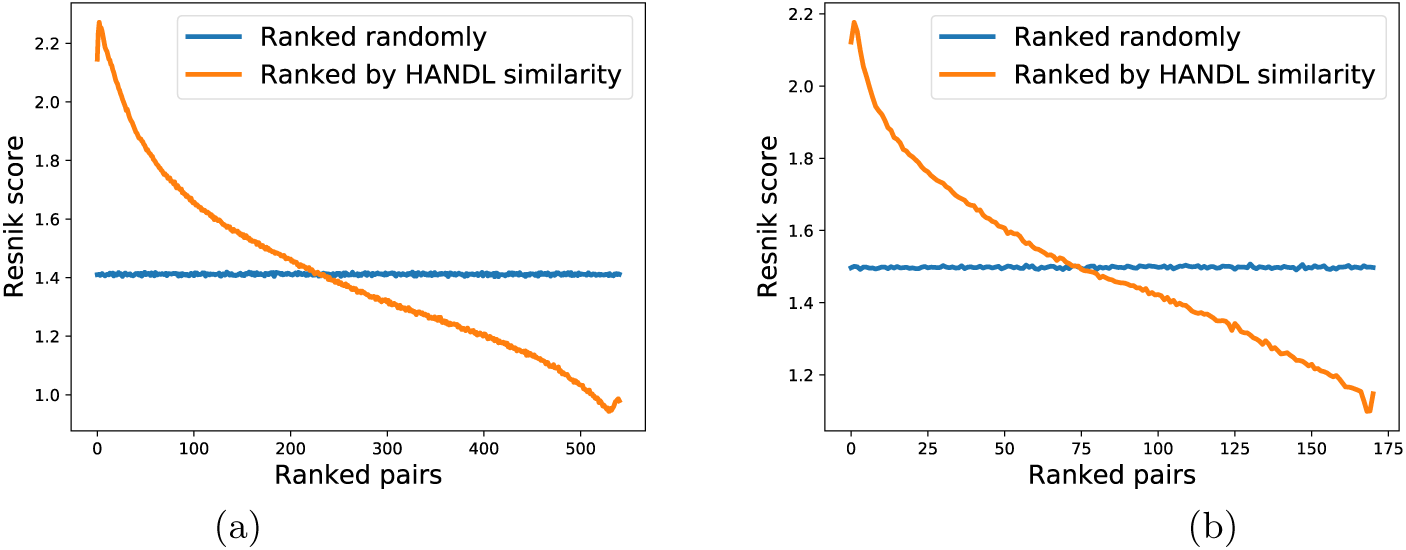
Relationship between cross-species Resnik similarity and Handl homology score, for (a) human (source) - yeast (target) and (b) mouse (source) - yeast (target) comparison.

**Fig. S2:**
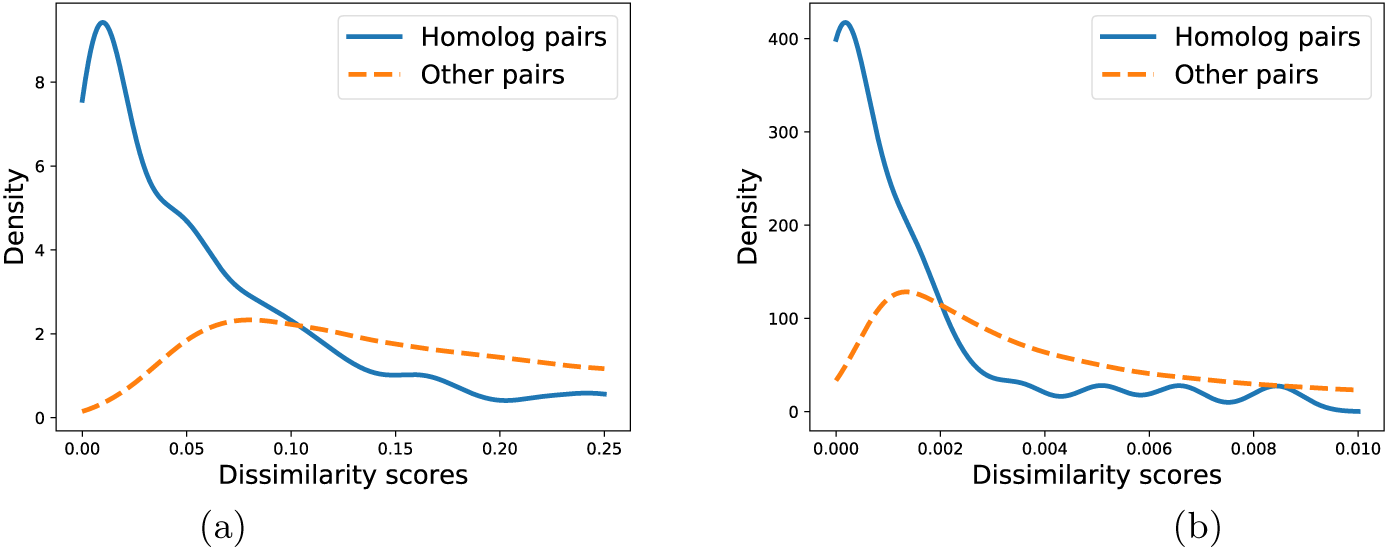
Distribution of Handl dissimilarity scores for homologs, compared to distribution for all protein pairs, for (a) human (source) - yeast (target) and (b) mouse (source) - yeast (target) comparison.

## References

1. Ahmet Aladag and Cesim Erten. Spinal: scalable protein interaction network alignment. Bioinformatics, 29(7):917–924, 2013.

2. Adrian M Altenhoff, Romain A Studer, Marc Robinson-Rechavi, and Christophe Dessimoz. Resolving the ortholog conjecture: orthologs tend to be weakly, but significantly, more similar in function than paralogs. PLoS Comput Biol, 8(5):e1002514, 2012.

3. Stephen F. Altschul, Warren Gish, Webb Miller, Eugene W. Myers, and David J. Lipman. Basic local alignment search tool. Journal of Molecular Biology, 215(3):403–410, 1990.

4. Michael Ashburner, Catherine A. Ball, Judith A. Blake, David Botstein, Heather Butler, et al. Gene ontology: tool for the unification of biology. Nature Genetics, 25(1):25–29, 2000.

5. Yoshua Bengio, Olivier Delalleau, and Nicolas Le Roux. Label propagation and quadratic criterion. In Olivier Chapelle, Bernhard Scholkopf, and Alexander Zien, editors, Semi-Supervised Learning, chapter 11. The MIT Press, 2006.

6. Charles Boone, Howard Bussey, and Brenda J Andrews. Exploring genetic interactions and networks with yeast. Nature Reviews Genetics, 8(6):437–449, 7 2007.

7. Kevin R Brown and Igor Jurisica. Unequal evolutionary conservation of human protein interactions in interologous networks. Genome biology, 8(5):R95, 2007.

8. Mengfei Cao, Hao Zhang, Jisoo Park, Noah Daniels, Mark Crovella, et al. Going the distance for protein function prediction: A new distance metric for protein interaction networks. PLOS One, 8(10):e76339, October 2013.

9. Andrew Chatraryamontri, Rose Oughtred, Lorrie Boucher, Jennifer Rust, Christie Chang, et al. The biogrid interaction database: 2017 update. Nucleic Acids Research, 45(D1):D369–D379, 2017.

10. JM Cherry, C Adler, C Ball, SA Chervitz, SS Dwight, ET Hester, Y Jia, G Juvik, T Roe, M Schroeder, S Weng, and D Botstein. SGD: Saccharomyces genome database. Nucleic Acids Res, 26:73–79, 1998.

11. Hyunghoon Cho, Bonnie Berger, and Jian Peng. Compact integration of multi-network topology for functional analysis of genes. Cell Systems, 3(6):540–548.e5, 2016.

12. Connor Clark and Jugal Kalita. A comparison of algorithms for the pairwise alignment of biological networks. Bioinformatics, 30(16):2351–2359, 2014.

13. Sean R Collins, Kyle M Miller, Nancy L Maas, Assen Roguev, Jeffrey Fillingham, et al. Functional dissection of protein complexes involved in yeast chromosome biology using a genetic interaction map. Nature, 446(7137):806–810, 7 2007.

14. Lenore Cowen, Trey Ideker, Benjamin J Raphael, and Roded Sharan. Network propagation: a universal amplifier of genetic associations. Nature Reviews Genetics, 2017.

15. Jesse Davis and Mark Goadrich. The relationship between precision-recall and roc curves. pages 233–240, 2006.

16. Raamesh Deshpande, Michael K Asiedu, Mitchell Klebig, Shari Sutor, Elena Kuzmin, et al. A comparative genomic approach for identifying synthetic lethal interactions in human cancer. Cancer Research, 73(20):6128–6136, 2013.

17. W M Fitch. Distinguishing homologous from analogous proteins. Systematic zoology, 19(2):99–113, 1970.

18. François Fouss, Kevin Francoisse, Luh Yen, Alain Pirotte, and Marco Saerens. An experimental investigation of kernels on graphs for collaborative recommendation and semisupervised classification. Neural Netw., 31:53–72, July 2012.

19. François Fouss, Marco Saerens, and Masashi Shimbo. Algorithms and Models for Network Data and Link Analysis. Cambridge University Press, 2016.

20. Andrea Franceschini, Damian Szklarczyk, Sune Frankild, Michael Kuhn, Milan Simonovic, Alexander Roth, Jianyi Lin, Pablo Minguez, Peer Bork, Christian von Mering, and Lars J. Jensen. String v9.1: protein-protein interaction networks, with increased coverage and integration. Nucleic Acids Research, 41(D1):D808–D815, 2013.

21. Adam Frost, Marc G Elgort, Onn Brandman, Clinton Ives, Sean R Collins, et al. Functional repurposing revealed by comparing s.pombe and s.cerevisiae genetic interactions. Cell, 149(6):1339–1352, 11 2012.

22. Jihun Ham, Daniel D Lee, and Lawrence K Saul. Semisupervised alignment of manifolds. In AISTATS, pages 120–127, 2005.

23. Hitomi Hasegawa and Liisa Holm. Advances and pitfalls of protein structural alignment. Current Opinion in Structural Biology, 19(3):341–348, 9 2009.

24. Somaye Hashemifar and Jinbo Xu. Hubalign: an accurate and efficient method for global alignment of protein–protein interaction networks. Bioinformatics, 30(17):i438–i444, 2014.

25. Sohyun Hwang, Eiru Kim, Sunmo Yang, Edward M Marcotte, and Insuk Lee. Morphin: a web tool for human disease research by projecting model organism biology onto a human integrated gene network. Nucleic Acids Research, 42(Web Server issue):W147–W153, 07 2014.

26. Alexandra Jacunski, Scott J. Dixon, and Nicholas P. Tatonetti. Connectivity homology enables inter-species network models of synthetic lethality. PLOS Computational Biology, 11(10):e1004506, 2015.

27. Lars J. Jensen, Michael Kuhn, Manuel Stark, Samuel Chaffron, Chris Creevey, Jean Muller, Tobias Doerks, Philippe Julien, Alexander Roth, Milan Simonovic, Peer Bork, and Christian von Mering. String 8–a global view on proteins and their functional interactions in 630 organisms. Nucleic acids research, 37(Database issue):D412–416, January 2009.

28. Ehsan Kazemi, Hamed Hassani, Matthias Grossglauser, and Hassan Pezeshgi Modarres. Proper: global protein interaction network alignment through percolation matching. BMC Bioinformatics, 17(1):527, Dec 2016.

29. Vikram Khurana, Jian Peng, Chee Chung, Pavan K Auluck, Saranna Fanning, et al. Genome-scale networks link neurodegenerative disease genes to -synuclein through specific molecular pathways. Cell Systems, 4(2):157–170.e14, 2017.

30. Gunnar W Klau. A new graph-based method for pairwise global network alignment. BMC bioinformatics, 10(Suppl 1):S59, 2009.

31. Risi Imre Kondor and John Lafferty. Diffusion kernels on graphs and other discrete input spaces. In ICML, volume 2, pages 315–322, 2002.

32. Eugene V Koonin, Natalie D Fedorova, John D Jackson, Aviva R Jacobs, Dmitri M Krylov, et al. A comprehensive evolutionary classification of proteins encoded in complete eukaryotic genomes. Genome Biology, 5(2):1–28, 3 2003.

33. G. R. Lanckriet, M. Deng, N. Cristianini, M. I. Jordan, and W. S. Noble. Kernel-based data fusion and its application to protein function prediction in yeast. In Pacific Symposium on Biocomputing, pages 300–311, 2004.

34. Christina S. Leslie, Eleazar Eskin, Adiel Cohen, Jason Weston, and William Stafford Noble. Mismatch string kernels for discriminative protein classification. Bioinformatics, 20(4):467, 2004.

35. Stanley Letovsky and Simon Kasif. Predicting protein function from protein/protein interaction data: a probabilistic approach. Bioinformatics, 19(suppl 1):i197–i204, 2003.

36. Chung-Shou Liao, Kanghao Lu, Michael Baym, Rohit Singh, and Bonnie Berger. Isorankn: spectral methods for global alignment of multiple protein networks. Bioinformatics, 25(12):i253–i258, 2009.

37. Nöel Malod-Dognin and Nataša Pržulj. L-graal: Lagrangian graphlet-based network aligner. Bioinformatics, page btv130, 2015.

38. Kriston L McGary, Tae Park, John O Woods, Hye Cha, John B Wallingford, and Edward M Marcotte. Systematic discovery of nonobvious human disease models through orthologous phenotypes. Proceedings of the National Academy of Sciences, 107(14):6544–6549, 10 2010.

39. Tijana Milenković, Weng Leong Ng, Wayne Hayes, and Nataša Pržulj. Optimal network alignment with graphlet degree vectors. Cancer Informatics, 9:121–137, 2010.

40. Florian L. Muller, Elisa A. Aquilanti, and Ronald A. DePinho. Collateral lethality: A new therapeutic strategy in oncology. Trends in Cancer, 1(3):161–173, 2015.

41. E. Nabieva, K. Jim, A. Agarwal, B. Chazelle, and M. Singh. Whole-proteome prediction of protein function via graph-theoretic analysis of interaction maps. Bioinformatics, 21(supplement 1):i302–i310, 2005.

42. Naoki Nariai. Probabilistic Integration of Heterogeneous, Contextual, and Cross-Species Genome-Wide Data for Protein Function Prediction. PhD thesis, Boston University, 2010.

43. M. E. J. Newman, S. H. Strogatz, and D. J. Watts. Random graphs with arbitrary degree distributions and their applications. Phys. Rev. E, 64:026118, Jul 2001.

44. Behnam Neyshabur, Ahmadreza Khadem, Somaye Hashemifar, and Seyed Shahriar Arab. Netal: a new graph-based method for global alignment of protein–protein interaction networks. Bioinformatics, 29(13):1654–1662, 2013.

45. Sebastian M.B. Nijman. Synthetic lethality: General principles, utility and detection using genetic screens in human cells. FEBS Letters, 585(1):1–6, 11 2011.

46. Nigel J O’Neil, Melanie L Bailey, and Philip Hieter. Synthetic lethality and cancer. Nature Reviews Genetics, 2017.

47. G Ostlund, T Schmitt, K Forslund, T Kostler, D N Messina, et al. Inparanoid 7: new algorithms and tools for eukaryotic orthology analysis. Nucleic Acids Research, 38(Database):D196–D203, 9 2009.

48. Sinno Jialin Pan and Qiang Yang. A survey on transfer learning. IEEE Trans. on Knowl. and Data Eng., 22(10):1345–1359, October 2010.

49. Yungki Park and Edward M Marcotte. Flaws in evaluation schemes for pair-input computational predictions. Nature Methods, 9(12):1134–1136, 12 2012.

50. Rob Patro and Carl Kingsford. Global network alignment using multiscale spectral signatures. Bioinformatics, 28(23):3105–3114, 2012.

51. Catia Pesquita, Daniel Faria, Andr O. Falco, Phillip Lord, and Francisco M. Couto. Semantic similarity in biomedical ontologies. PLoS Comput Biol, 5(7):1–12, 07 2009.

52. Mark E Peterson, Feng Chen, Jeffery G Saven, David S Roos, Patricia C Babbitt, and Andrej Sali. Evolutionary constraints on structural similarity in orthologs and paralogs. Protein Science, 18(6):1306–1315, 2009.

53. Hang TT Phan and Michael JE Sternberg. Pinalog: a novel approach to align protein interaction networks: implications for complex detection and function prediction. Bioinformatics, 28(9):1239–1245, 2012.

54. Philip Resnik. Semantic similarity in a taxonomy: An information-based measure and its application to problems of ambiguity in natural language. J. Artif. Intell. Res.(JAIR), 11:95–130, 1999.

55. Igor B Rogozin, David Managadze, Svetlana A Shabalina, and Eugene V Koonin. Gene family level comparative analysis of gene expression in mammals validates the ortholog conjecture. Genome biology and evolution, 6(4):754–762, 2014.

56. Assen Roguev, Sourav Bandyopadhyay, Martin Zofall, Ke Zhang, Tamas Fischer, et al. Conservation and rewiring of functional modules revealed by an epistasis map in fission yeast. Science, 322(5900):405–410, 8 2008.

57. Eric W Sayers, Tanya Barrett, Dennis A Benson, Evan Bolton, Stephen H Bryant, et al. Database resources of the national center for biotechnology information. Nucleic acids research, 39(suppl 1):D38–D51, 2011.

58. Rohit Singh, Jinbo Xu, and Bonnie Berger. Global alignment of multiple protein interaction networks with application to functional orthology detection. Proceedings of the National Academy of Sciences, 105(35):12763–12768, 2008.

59. Jimin Song and Mona Singh. How and when should interactome-derived clusters be used to predict functional modules and protein function? Bioinformatics, 25(23):3143–3150, 2009.

60. C. Stark, B.J. Breitkreutz, T. Reguly, L. Boucher, A. Breitkreutz, and M. Tyers. BioGRID: a general repository for interaction datasets. Nucleic Acids Res, 34(Database Issue):D535–D539, 2006.

61. Baochen Sun, Jiashi Feng, and Kate Saenko. Return of frustratingly easy domain adaptation. In Proceedings of the Thirtieth AAAI Conference on Artificial Intelligence, AAAI’16, pages 2058–2065. AAAI Press, 2016.

62. Mark G. F. Sun, Martin Sikora, Michael Costanzo, Charles Boone, and Philip M. Kim. Network evolution: Rewiring and signatures of conservation in signaling. PLoS Computational Biology, 8(3):e1002411, 12 2012.

63. The UniProt Consortium. Uniprot: the universal protein knowledgebase. Nucleic Acids Research, 45(D1):D158–D169, 2017.

64. Fabio Vandin, Eli Upfal, and Benjamin J. Raphael. Algorithms for detecting significantly mutated pathways in cancer. Journal of Computational Biology, 18(3):507–522, 2016/11/07 2011.

65. Vipin Vijayan and Tijana Milenkovic. Multiple network alignment via multimagna++. arXiv preprint arXiv:1604.01740, 2016.

66. Chang Wang and Sridhar Mahadevan. Manifold alignment using procrustes analysis. In Proceedings of the 25th International Conference on Machine Learning, pages 1120–1127, 2008.

67. D. Zhou and B. Schölkopf. A regularization framework for learning from graph data. In ICML Workshop on Statistical Relational Learning and Its Connections to Other Fields, pages 132–137, 2004.

